# Modelling and simulation of the interactions between the cardiovascular system, VA ECMO and IABP: when is it an appropriate combination?

**DOI:** 10.1101/2025.02.03.636021

**Authors:** Beatrice De Lazzari, Massimo Capoccia, Roberto Badagliacca, Marc O Maybauer, Claudio De Lazzari

## Abstract

Veno-arterial extracorporeal membrane oxygenation (VA ECMO) for the management of refractory cardiogenic shock (CS) has been widely used in recent years. Increased left ventricular (LV) afterload induced by retrograde flow remains a limiting factor, which is particularly evident during peripheral VA ECMO support. The concomitant use of the intra-aortic balloon pump (IABP) is an established strategy to achieve LV unloading during VA ECMO support. Nevertheless, there remains controversy about the combined use of IABP during central or peripheral VA ECMO in terms of beneficial effects and outcome. We developed a simulation setting to study left ventricular unloading with IABP during peripheral and central VA ECMO using CARDIOSIM^©^, an established software simulator of the cardiovascular system. The aim was to quantitatively evaluate potential differences between the two VA ECMO configurations and ascertain the true beneficial effects compared to VA ECMO alone.

## Introduction

The use of veno-arterial extracorporeal membrane oxygenation (VA ECMO) for the management of refractory cardiogenic shock (CS) has significantly increased in recent years (Meani, 2017). Despite the established benefit of VA ECMO support, increased left ventricular (LV) afterload induced by retrograde flow remains a limiting factor which is particularly evident during peripheral VA ECMO insertion. Retrograde flow can significantly limit LV ejection leading to blood stasis and LV distension. Reduced or absent forward flow may also develop due to a mismatch between LV afterload, preload and contractility. There is increasing evidence that LV unloading during VA ECMO support is beneficial (Lorusso, 2022). The concomitant use of the intra-aortic balloon pump (IABP) is an established strategy for this purpose in view of its wide availability, ease of insertion and maintenance in the cardiac intensive care unit. Patients managed with the combined use of VA ECMO and IABP demonstrated significant lower in-hospital and 30-day mortality compared to those managed with VA ECMO alone without an increase in complications (Nishi, 2022; Li, 2019; Vallabhajosyula, 2018). This matter has not been free of controversy. Despite increasing evidence confirming the beneficial effect of LV unloading on VA ECMO, there have been reports questioning its translation into better outcomes (Björnsdóttir, 2022; Hasde, 2021; Lin, 2016; Wang, 2020; Tepper, 2019). Nevertheless, the evidence supporting the combined use of VA ECMO and IABP in postcardiotomy failure and cardiogenic shock remains quite overwhelming (Baran, 2024; Wang, 2024; Chen, 2019; Kida, 2022; LoForte, 2016; Nuding and Werdan, 2017; Zeng, 2022). VA ECMO support can be achieved either peripherally or centrally according to the needs and the context with some evidence supporting both approaches (Bréchot, 2018; Raffa, 2019; Xu, 2022) although the combined use of peripheral VA ECMO and IABP has been questioned (Tay, 2016; Yang, 2014). The effect of IABP on LV afterload and organ perfusion is complex and changes according to the degree of left ventricular systolic dysfunction. The crux of the balloon lies with the retrograde flow of peripheral VA ECMO. In the presence of severe left ventricular systolic dysfunction, the afterload may paradoxically increase during systole due to balloon deflation or “de-clamping” effect of the descending thoracic aorta. Balloon occlusion of the aorta in diastole may reduce VA ECMO-driven blood flow to the aortic root and arch attenuating myocardial and cerebral blood flow (Bělohlávek, 2012). When the degree of LV systolic dysfunction is less, lower VA ECMO support is required and the impact of retrograde flow affecting the intended effect of IABP is less. The resultant pseudo-pulsatile flow induced by IABP would improve regional microcirculation (Jung, 2009; Petroni, 2014). The addition of IABP to central VA ECMO leads to the reduction of intracavitary cardiac pressure and work with potential for myocardial recovery in a pig model (Djordjevic, 2022). A mock circulation loop study of IABP and concomitant ECMO support during different degree of LV failure suggests a strong relationship between IABP efficacy and ventricular contractility inviting to careful selection when considering concomitant VA ECMO and IABP support (Farag, 2022). Another experimental setting in a pig model has found higher coronary blood flow in central VA ECMO compared to peripheral VA ECMO regardless of flow conditions in the ascending aorta although the addition of IABP did not result in improved coronary blood flow but showed partial negative effect on the coronary arterial endothelial function (Gerfer, 2023). Computational studies have focused on wall shear stress and modulation of pulsatility induced by IABP (Gu, 2016; Gu, 2018; Gu, 2022). An in-vitro study based on a silicon phantom model of the circulation has shown increased coronary blood flow on VA ECMO support with further improvement following IABP insertion particularly with reduced heart rate (Reymond, 2021). Given these considerations, we developed a simulation setting to study left ventricular unloading with IABP during peripheral and central VA ECMO. The aim was to quantitatively evaluate potential differences between the two VA ECMO configurations and ascertain the true beneficial effects compared to VA ECMO alone.

## Material and Methods

In this study we assembled CARDIOSIM^©^ (De Lazzari, 2022; De Lazzari 2022) software platform as shown in Fig.1 The numerical model reproducing the behaviour of both ventricles, atria and septum has been previously described (De Lazzari, 2012; Capoccia, 2018). The numerical models of ascending and descending aorta with aortic arch, descending thoracic aorta (with thoracic resistance RTHOR), upper limbs and head, superior and inferior vena cava, renal and hepatic, splanchnic, abdominal and lower limbs circulation have also been described in detail (De Lazzari 2021).

**Figure 1.**
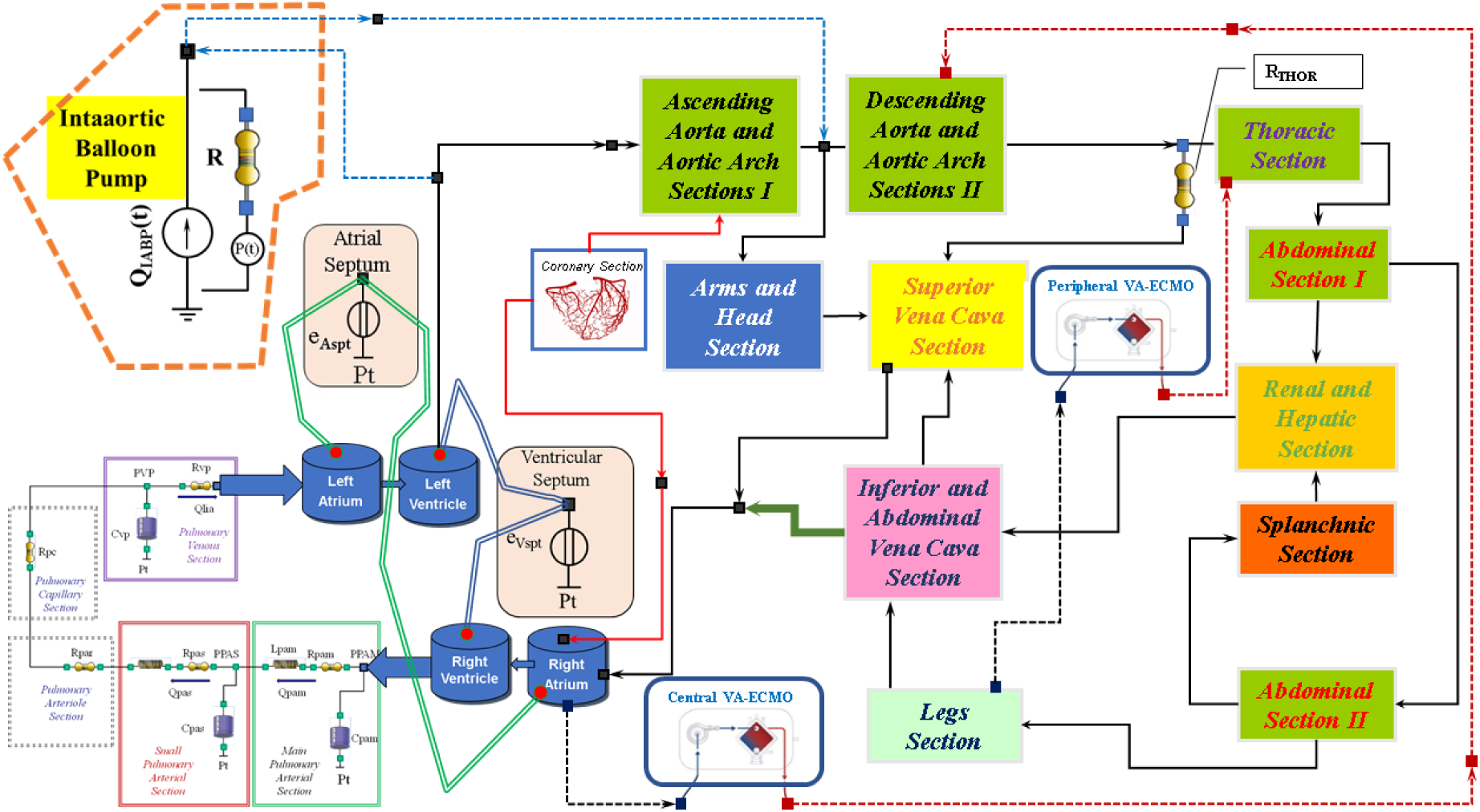
Electric analogue of the cardiovascular system with intra-aortic balloon pump (IABP), central and peripheral VA ECMO. The different sections of ascending and descending aorta with aortic arch, descending thoracic aorta (with thoracic resistance RTHOR), upper limbs and head, superior and inferior vena cava, renal and hepatic, splanchnic, abdominal and lower limbs circulation have been modelled with resistance, inertance and compliance (RLC) elements (De Lazzari 2021). The coronary circulation was modelled with RC elements. The numerical model reproducing central and peripheral VA ECMO configurations have been previously described (De Lazzari 2021).

The coronary circulation was modelled using resistance and compliance (RC) elements (De Lazzari, 2007). The network between the right ventricle and the left atria was reproduced using resistance, inertance and compliance (RLC) elements (Capoccia 2018).

The numerical model reproducing central and peripheral VA ECMO configurations (Figures 1 and 2) have been previously described (De Lazzari 2021). Central VA ECMO draws blood from the right atrium (RA) and ejects it into the ascending aorta (Asc Ao). Peripheral VA ECMO draws blood from the femoral vein (FV) and ejects it into the femoral artery (FA). For the purposes of the simulations, the two circuits have been configured as shown in Figure 2.

**Figure 2.**
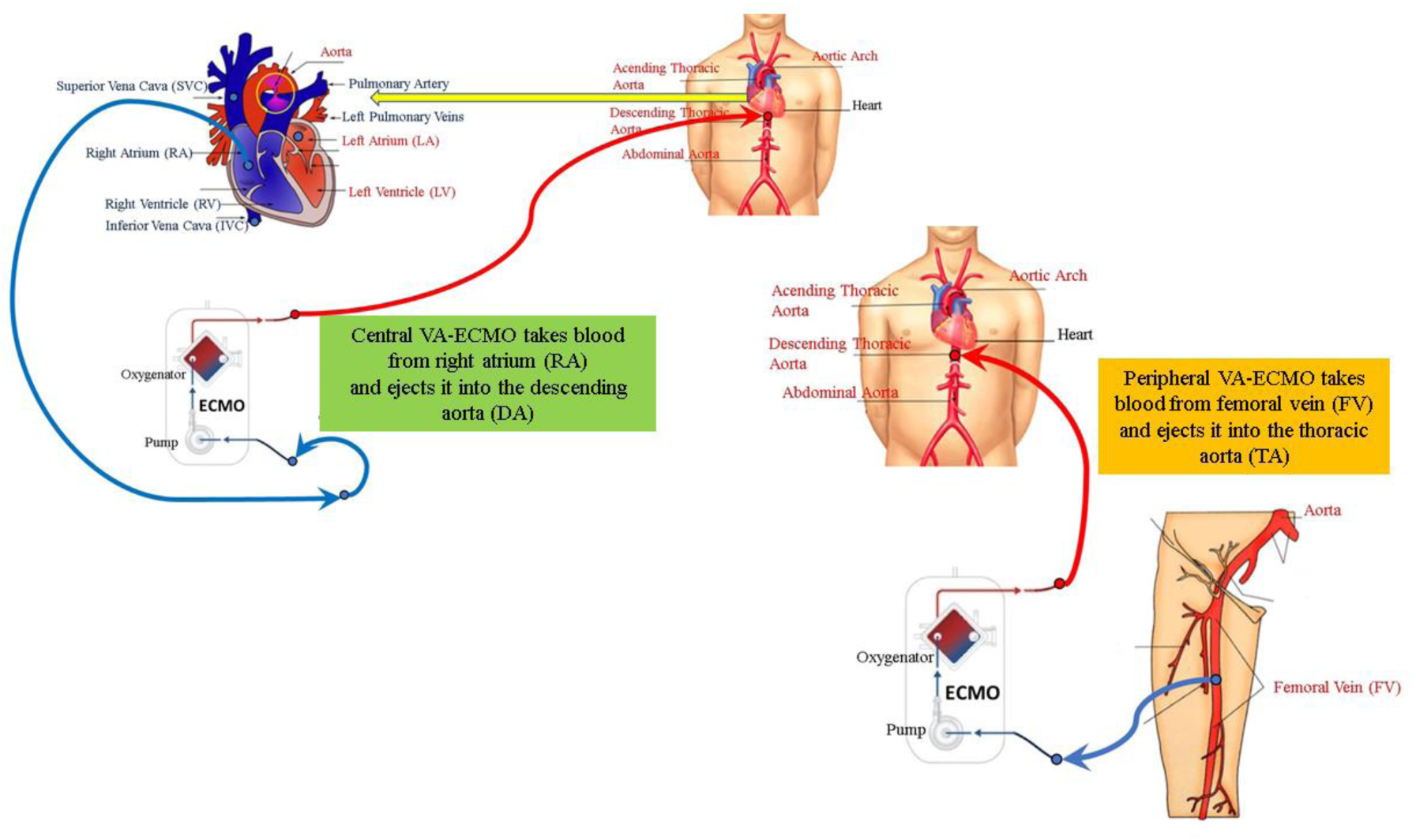
Schematic representation of central and peripheral VA ECMO. The circuits have been slightly modified for the purposes of the simulations. Central VA ECMO draws blood from the right atrium (RA) and ejects it into the descending aorta (DA). Peripheral VA ECMO draws blood from the femoral vein (FV) and ejects it into the thoracic aorta (TA).

The IABP model, inserted in the arterial tree, is considered a flow source Q_IABP_(t) as follows: during diastole the balloon inflates generating positive flow whilst during systole the balloon deflates leading to negative flow. The flow source Q_IABP_(t) may be replaced by a pneumatic pressure source P(t), representing the compressed gas reservoir, and by resistance (R) representing the total gas delivery resistance of the system (Figure 1). The pneumatic source P(t) has been modelled describing the ejection and the filling phase separately as the air outflow from a high-pressure tank, connected to the pressure source, and the air outflow from a lower-pressure tank connected to the vacuum source (De Lazzari 2023).

In the first step of the study, a cardiogenic shock (CS) patient was reproduced in accordance with guidelines using the network configuration shown in Figure 1. The software simulator parameters were adjusted to obtain a systolic blood pressure < 90 mmHg, a systemic vascular resistance index (SVRi) < 1800 (dynes/sec/m^2^/cm^−5^), pulmonary capillary wedge pressure (PCWP) > 18 mmHg and a cardiac index (CI) < 1.8 L/min/m^2^. In the second step, VA ECMO support was activated as central and peripheral configuration. In both conditions, the simulations were performed setting the pump rotational speed to 3000, 3500 and 4000 rpm, respectively. Finally, VA ECMO connected both centrally and peripherally with the same pump rotational speed was switched on in conjunction with IABP. Intra-aortic balloon counterpulsation was set with a drive pressure of 260 mmHg and vacuum pressure of -10 mmHg. We investigated how the following haemodynamic variables were affected by central and peripheral VA ECMO, both with and without IABP support:

- minimum, maximum and mean values of left atrial pressure (LAP), aortic pressure (AoP) and pulmonary arterial pressure (PAP),
- pulmonary capillary wedge pressure (PCWP),
- total flow (native plus assistance flow),
- coronary blood flow (CBF),
- cardiac index (CI),
- left ventricular end systolic (diastolic) volume LVESV (LVEDV),
- right ventricular end systolic (diastolic) volume RVESV (RVEDV),
- systemic vascular resistance index (SVRi).

In addition, the following energetic variables were investigated:

✓ left (right) atrial pressure-volume loop area LA-PVLA (RA-PVLA),
✓ left (right) ventricular external work LV-EW (RV-EW),
✓ left (right) ventricular pressure-volume area LV-PVA (RV-PVA).

Data analysis included the assessment of the percentage variations of the assistance obtained under central and peripheral VA ECMO with or without IABP support, estimated in comparison to pathological conditions.

## Results

Figure 3 shows the left and right ventricular pressure-volume loops (top panel) and the left atrial pressure-volume loop (bottom panel) of pathological conditions (dashed blue loops) and assisted conditions with central (red loops) and peripheral (black loops) VA ECMO support when the pump rotational speed was set to 3000 rpm. The figure’s bar graph gives the percentage changes of LVESV, LVEDV, RVESV, RVEDV and left atrial end systolic (diastolic) volume LAESV (LAEDV) compared to pathological conditions during the two methods of assistance. When the pump rotational speed was set to 3000 rpm, peripheral VA ECMO increased LVESV, LVEDV, RVESV, RVEDV LAESV, and LAEDV by more than 14%, whereas central VA ECMO increased LVESV (≅ 14%), LVEDV (≅ 6%) and LAESV (≅ 10%), but decreased RVESV and LAEDV by 4% and RVEDV by 8%.

**Figure 3.**
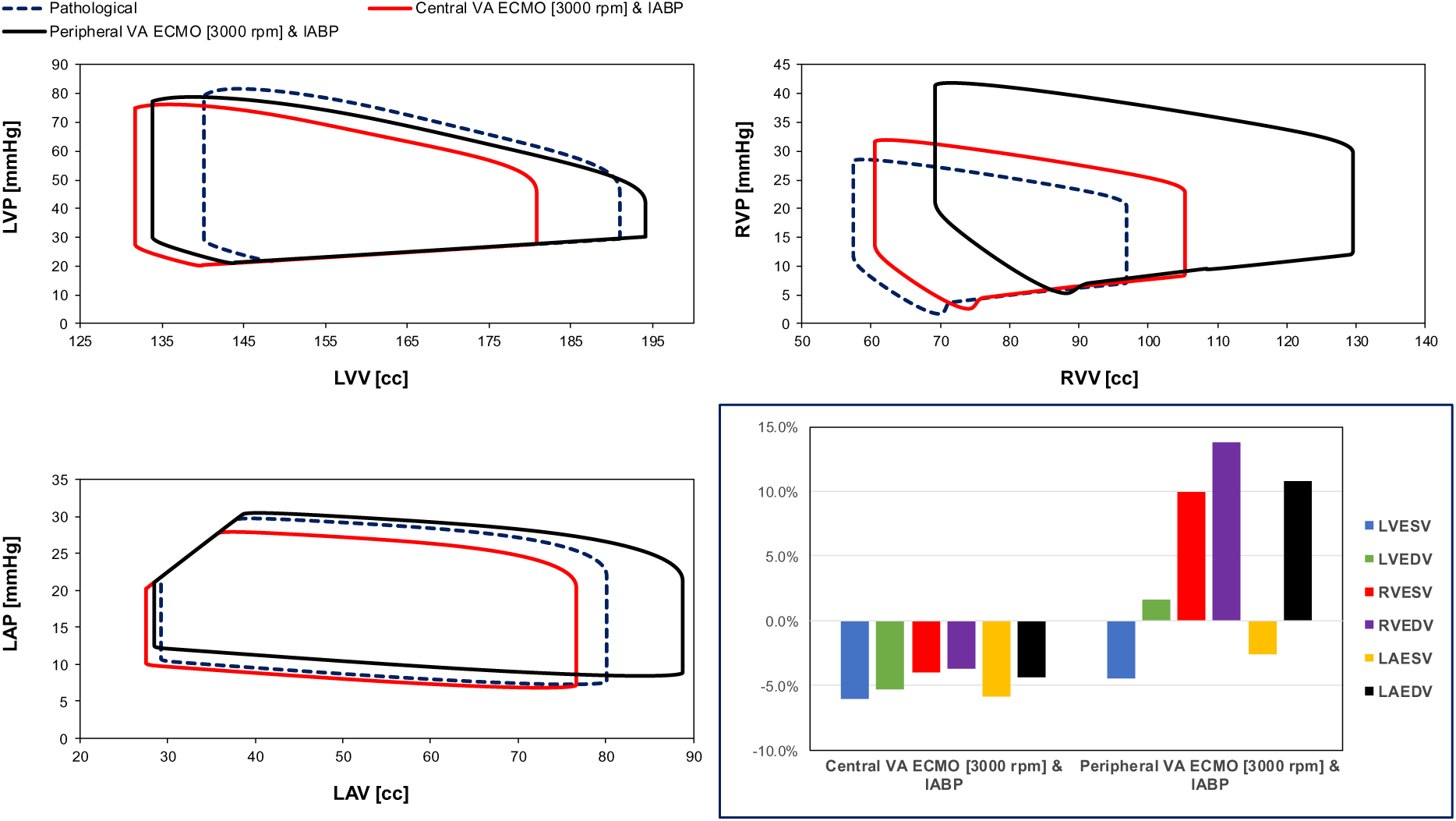
Top left (right) panel shows the left (right) ventricular pressure-volume loops reproduced by CARDIOSIM© to simulate the pathological conditions (dashed blue line) and the assisted conditions with central (red line) and peripheral (black line) VA ECMO. The pump rotational speed was set to 3000 rpm. The bottom left panel shows the left atrial pressure-volume loops. The bar graph (bottom right panel) shows the percentage changes of assisted conditions compared to pathological conditions in terms of LVESV, LVEDV, RVESV, RVEDV, LAESV and LAEDV.

Figure 4 reports the effects on the left and right ventricular pressure-volume loops as well as the left atrial pressure-volume loop induced by the simultaneous activation of VA ECMO and IABP. The pathological left (right) ventricular pressure-volume loop (dashed blue line), the assisted central VA ECMO with IABP left (right) ventricular pressure-volume loop (red line) and the assisted peripheral VA ECMO with IABP left (right) ventricular pressure-volume loop (black line) are all presented in the top left (right) panel. The left atrial pressure-volume loop of pathological conditions (dashed blue loop), the assisted conditions with central (red loop) and peripheral (black loop) VA ECMO both in conjunction with IABP support are available in the left bottom panel. When IABP was applied in conjunction with central VA ECMO support, LVESV, LVEDV, RVESV, RVEDV, LAESV, and LAEDV decreased, whereas peripheral VA ECMO and IABP support would decrease LVESV and LAESV but increase LVEDV, RVESV, RVEDV and LAEDV compared to pathological conditions.

**Figure 4.**
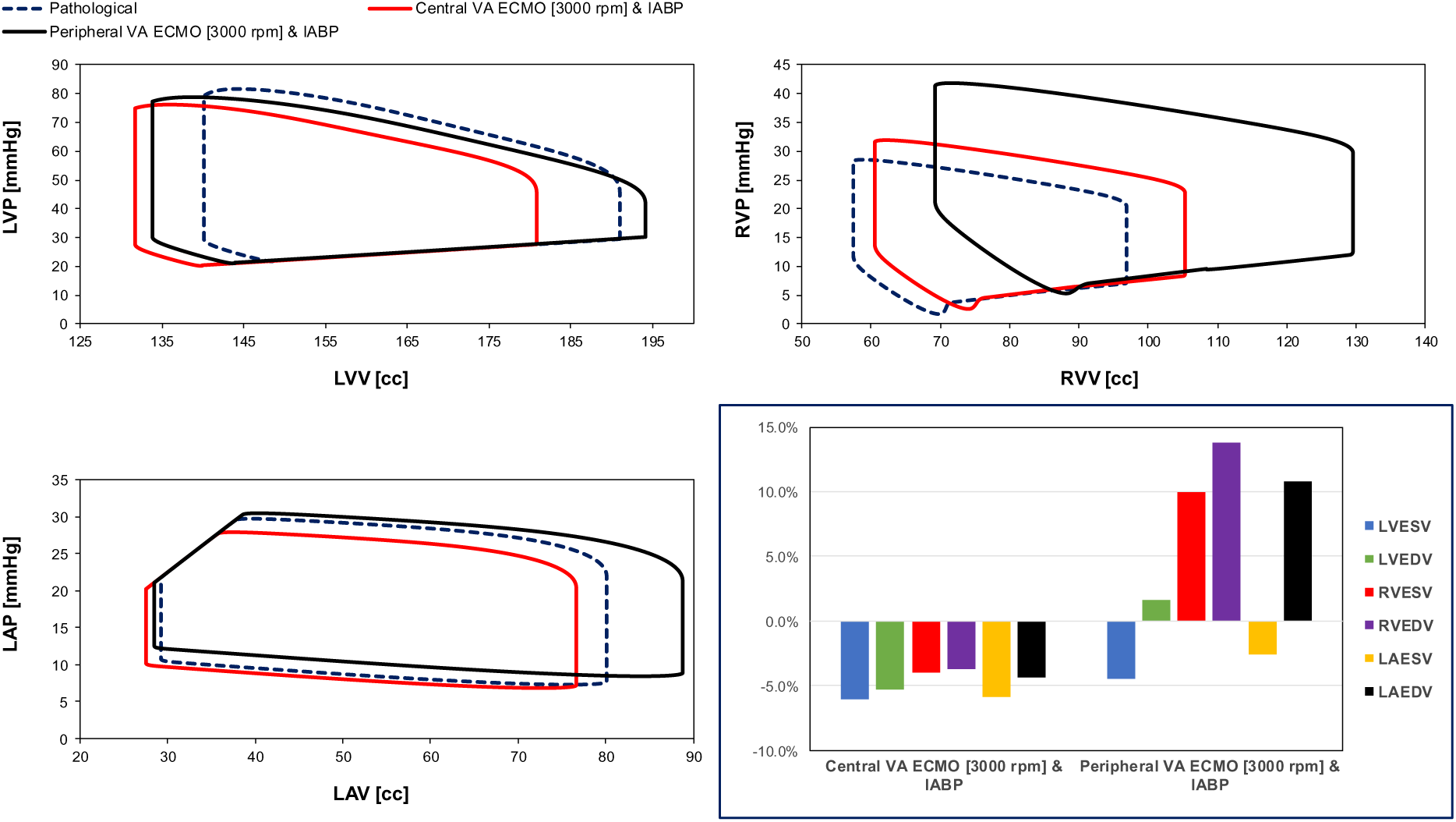
Top left (right) panel shows the left (right) ventricular pressure-volume loops reproduced by CARDIOSIM^©^ to simulate the pathological conditions (dashed blue line) and the assisted conditions with IABP in conjunction with central (red line) and peripheral (black line) VA ECMO, respectively. The pump rotational speed was set to 3000 rpm and the drive and vacuum IABP pressures were set to 260 mmHg and -10 mmHg, respectively. The bottom left panel shows the left atrial pressure-volume loops. The bar graph (bottom right panel) gives the percentage changes of assisted conditions compared to pathological conditions in terms of LVESV, LVEDV, RVESV, RVEDV, LAESV and LAEDV.

Concomitant central VA ECMO (with pump rotational speed set to 3000 rpm) and IABP support reduced both left ventricular and left atrial ESVs and EDVs.

When central VA ECMO was applied with pump rotational speeds of 3000 and 3500 rpm, with or without IABP assistance, SVRi increased from 43.4% to 83.83% (top graph in Figure 5), while no effect was seen with peripheral VA ECMO support. The same bar graph shows a decrease in cardiac index with central VA ECMO support (for every rotational speed), even after concomitant IABP activation. When peripheral VA ECMO was active, CI increased up to 44.86% (for 3500 rpm and activated IABP). Systemic elastance (Ea) and the ratio between systemic elastance and systemic arterial elastance (Eas) decreased when peripheral VA ECMO and IABP were both activated; a negligible effect was observed in the absence of IABP support (bottom graph in Figure 5). Pulmonary elastance was negligible following both types of assistance. When central VA ECMO was used, percentage changes in Ea/Ees ranging from 27.31% to 52.42% (for a 3000 rpm rotational pump speed) and percentage changes in systemic Ea ranging from 27.04% to 40.88% (for a 3500 rpm rotational pump speed) were observed when IABP support was turned off. The percentage changes in terms of total blood flow increased when the rotating pump speed of VA ECMO increased in both peripheral and central configurations (top graph in Figure 6). IABP support produced a 15% increase in Ea and Ea/Ees during central VA ECMO and a 20% decrease during peripheral VA ECMO.

**Figure 5.**
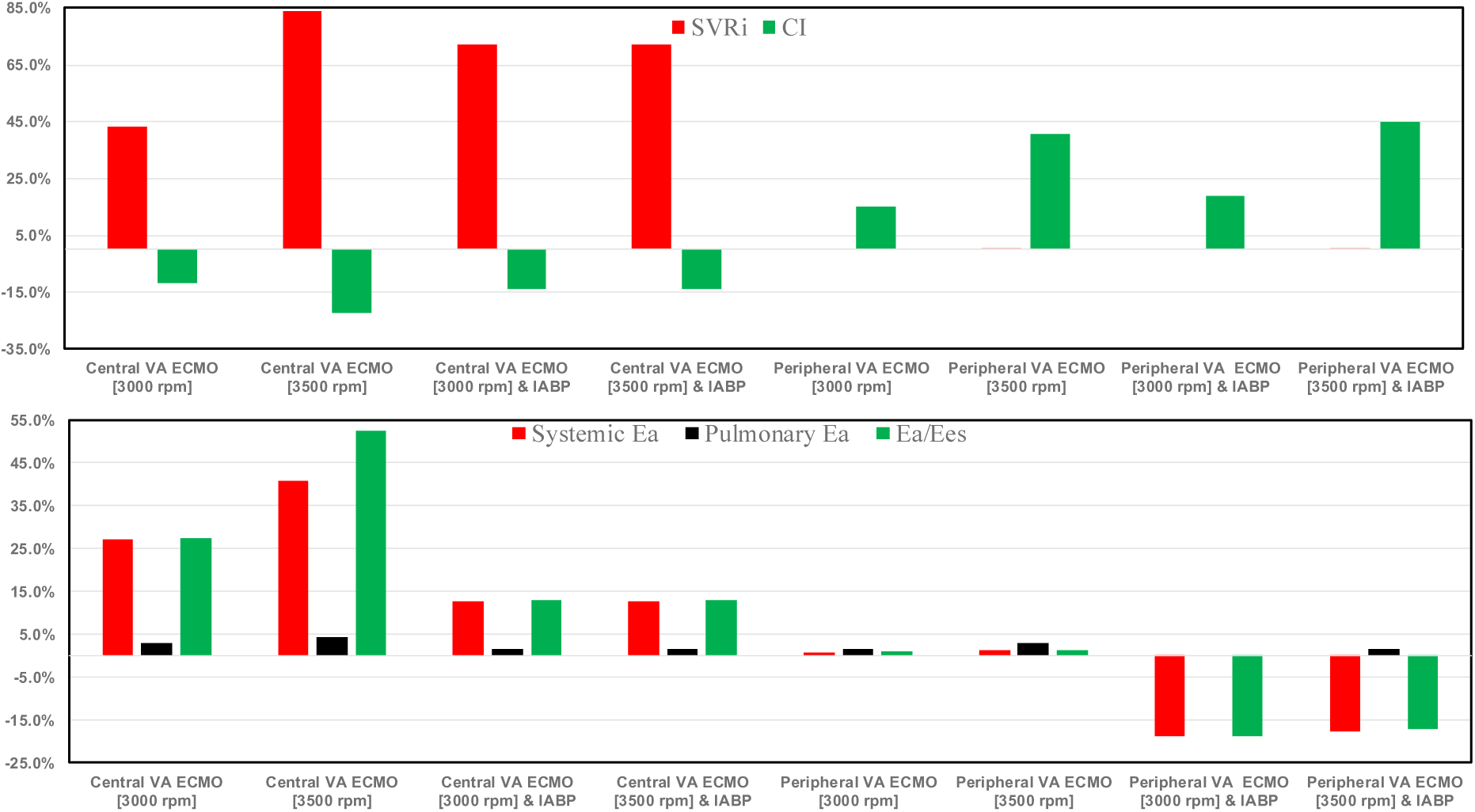
Top bar graph shows the percentage changes of SVRi and cardiac index in assisted conditions with central and peripheral VA ECMO driven at 3000 and 3500 rpm, as well as with and without IABP support, in relation to pathological conditions. The bottom bar graph shows the percentage changes of systemic and pulmonary Ea and Ea/Ees in relation to pathological conditions.

**Figure 6.**
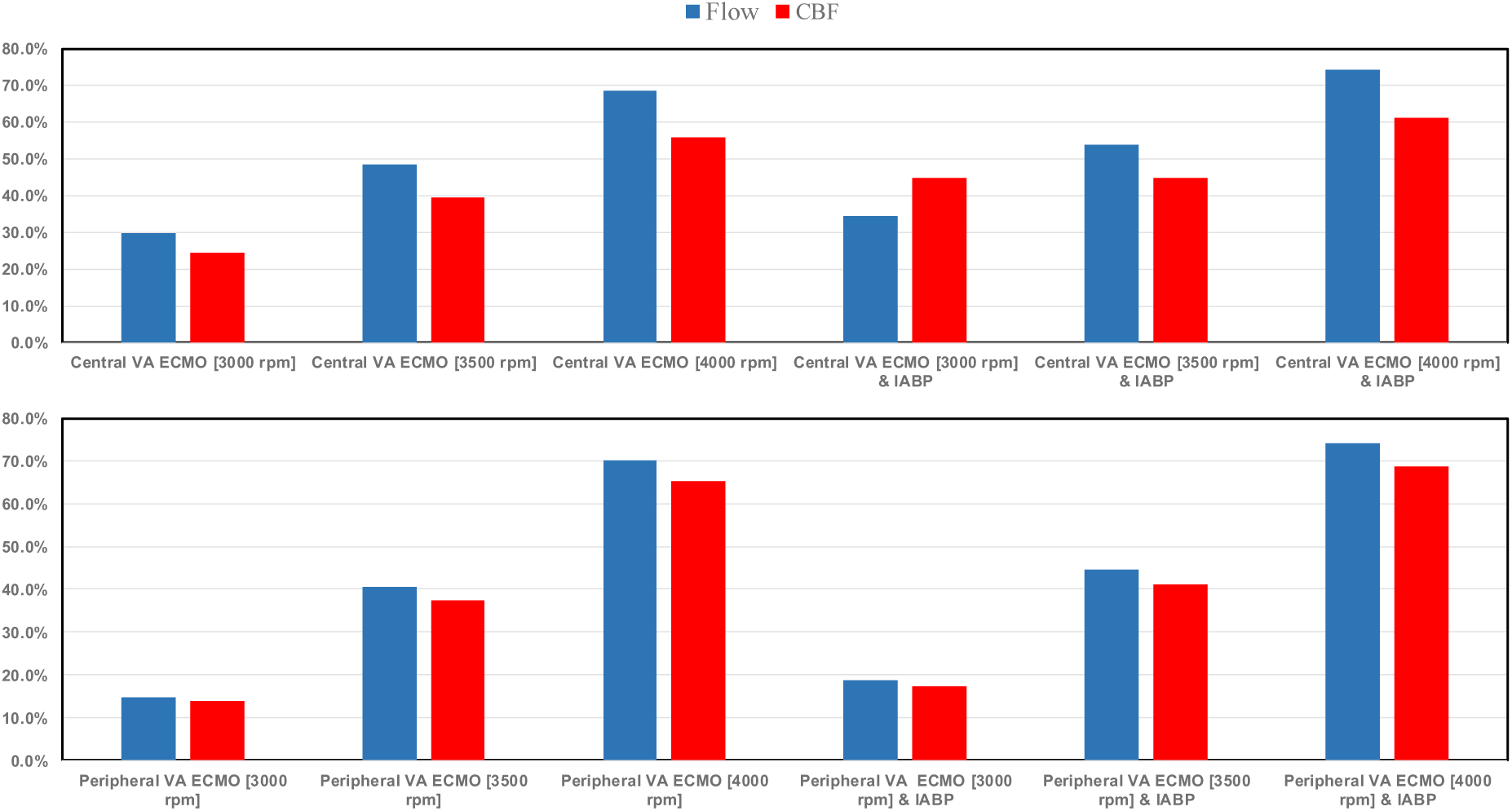
The top (bottom) bar graph shows the percentage changes of blood flow and coronary blood flow (CBF) in assisted conditions with central (peripheral) VA ECMO driven at 3000, 3500, and 4000 rpm, as well as with and without IABP support, in relation to pathological conditions.

The percentage changes in CBF when peripheral (central) VA ECMO was applied ranged between 65.15% and 68.66% (55.82% and 61.30%) when the pump rotational speed was set to 4000 rpm either without or with IABP, respectively (Figure 6). For central (peripheral) VA ECMO, CBF increased from 24.32% (13.78%) to 44.69% (17.21%) when the pump rotational speed was set to 3000 rpm, again without and with IABP.

Figure 7 shows the percentage changes in cerebral blood flow under assisted conditions with central and peripheral VA ECMO, as well as with and without IABP support, related to pathological conditions. The cerebral perfusion in both VA ECMO configurations increased as the pump rotational speed increased. IABP support did not seem to affect cerebral perfusion significantly.

**Figure 7.**
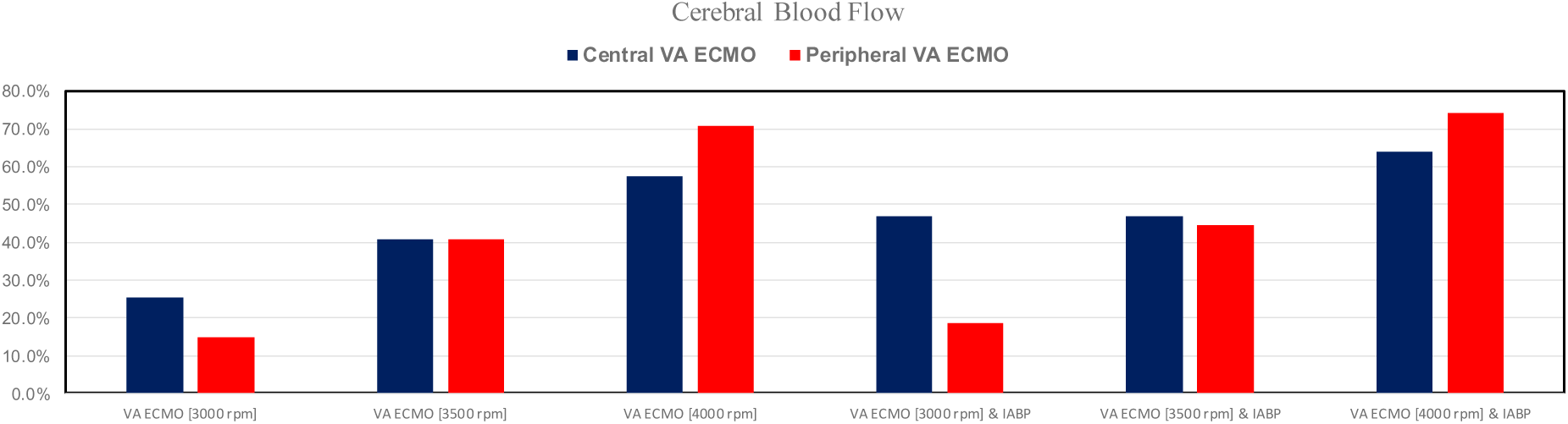
Percentage changes of cerebral blood flow under assisted conditions with central and peripheral VA ECMO driven at 3000, 3500, and 4000 rpm, as well as with and without IABP support, in relation to pathological conditions.

Figure 8 shows the percentage changes in the minimum values (panel A) of LAP, LVP, and AoP when central and peripheral VA ECMO was used with pump rotational speed set to 3000, 3500, and 4000 rpm with or without IABP. The minimum AoP value increased up to 64.62% (4000 rpm) when VA ECMO was connected centrally and the pump rotational speed was set to 3000, 3500, and 4000 rpm; IABP activation significantly reduced the minimum AoP values (panel A). Similar trends were observed when VA ECMO was connected peripherally either with or without IABP.

**Figure 8.**
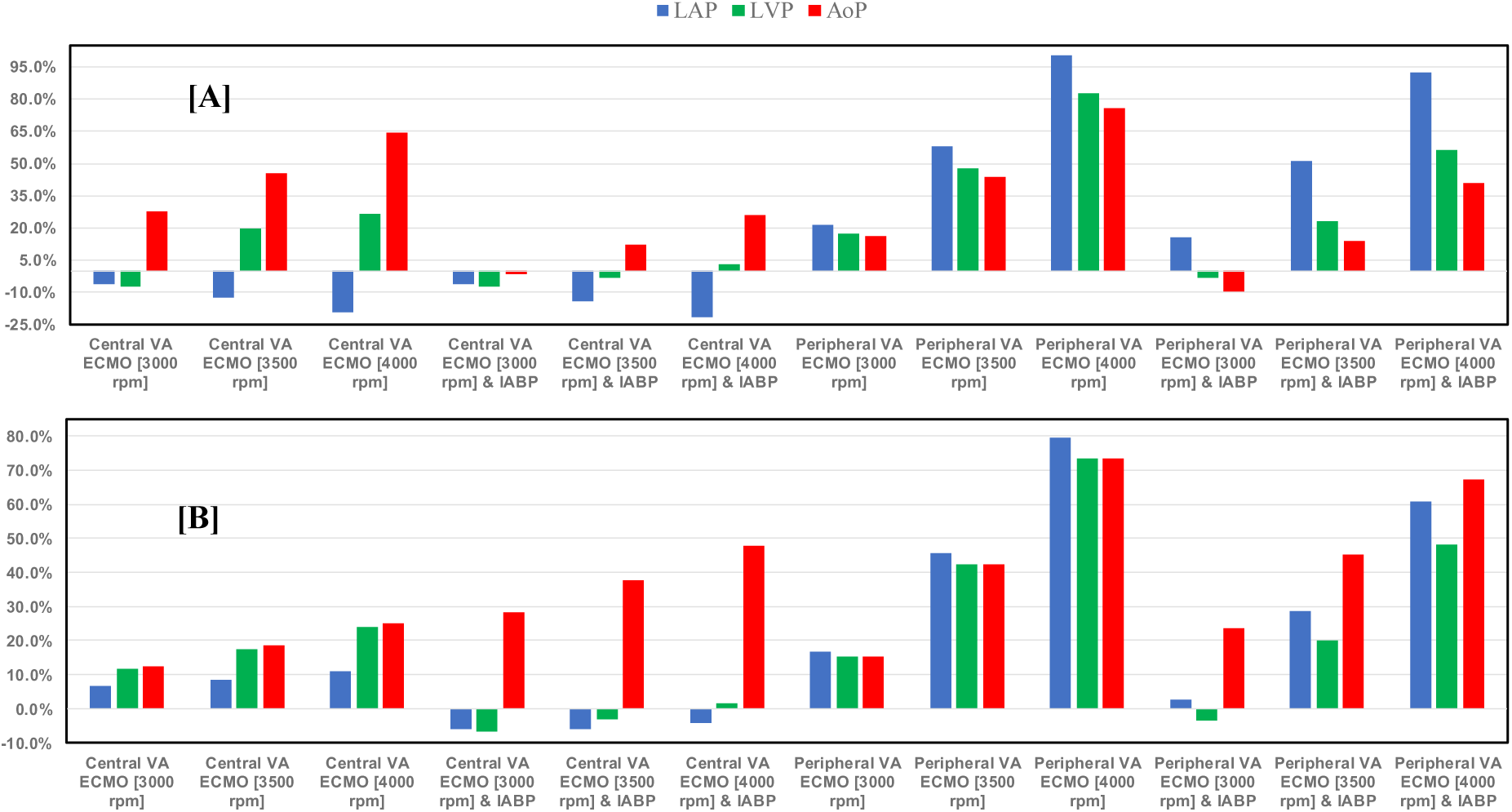
The top bar graph (A) shows the percentage changes in minimum value of LAP (wedge), left ventricular pressure (LVP) and AoP in assisted conditions with central/peripheral VA ECMO driven at 3000, 3500, and 4000 rpm, as well as with and without IABP support, in relation to pathological conditions. Bar graph (B) shows percentage changes in maximum LAP, LVP and AoP for central/peripheral VA ECMO with and without IABP support, in relation to pathological conditions.

Even with active IABP, the minimum LAP value fell when VA ECMO was connected centrally, but the minimum LAP and LVP values increased for peripheral VA ECMO.

Panel B (Figure 8) shows that when central VA ECMO pump rotational speed increased, the maximal AoP values also increased. This effect was more evident when IABP support was switched on. Peripheral VA ECMO raised LAP, LVP, and AoP maximum values above 73.5% (4000 rpm). Nevertheless, this effect decreased when IABP and peripheral VA ECMO were activated simultaneously. The maximum LAP and LVP values were comparable to those in non assisted conditions when IABP and central VA ECMO were activated simultaneously (for all pump rotational speeds) and when peripheral VA ECMO (3000 rpm) and IABP worked concomitantly.

Figure 9 shows the percentage changes in mean AoP value calculated in relation to pathological conditions for both central and peripheral VA ECMO assistance (with pump rotational speed set to 3000, 3500, and 4000 rpm), with and without IABP support. The mean aortic pressure values increased in each condition when IABP assistance was present; the greatest increase occurred with peripheral VA ECMO when the pump speed was set to 4000 rpm.

**Figure 9.**
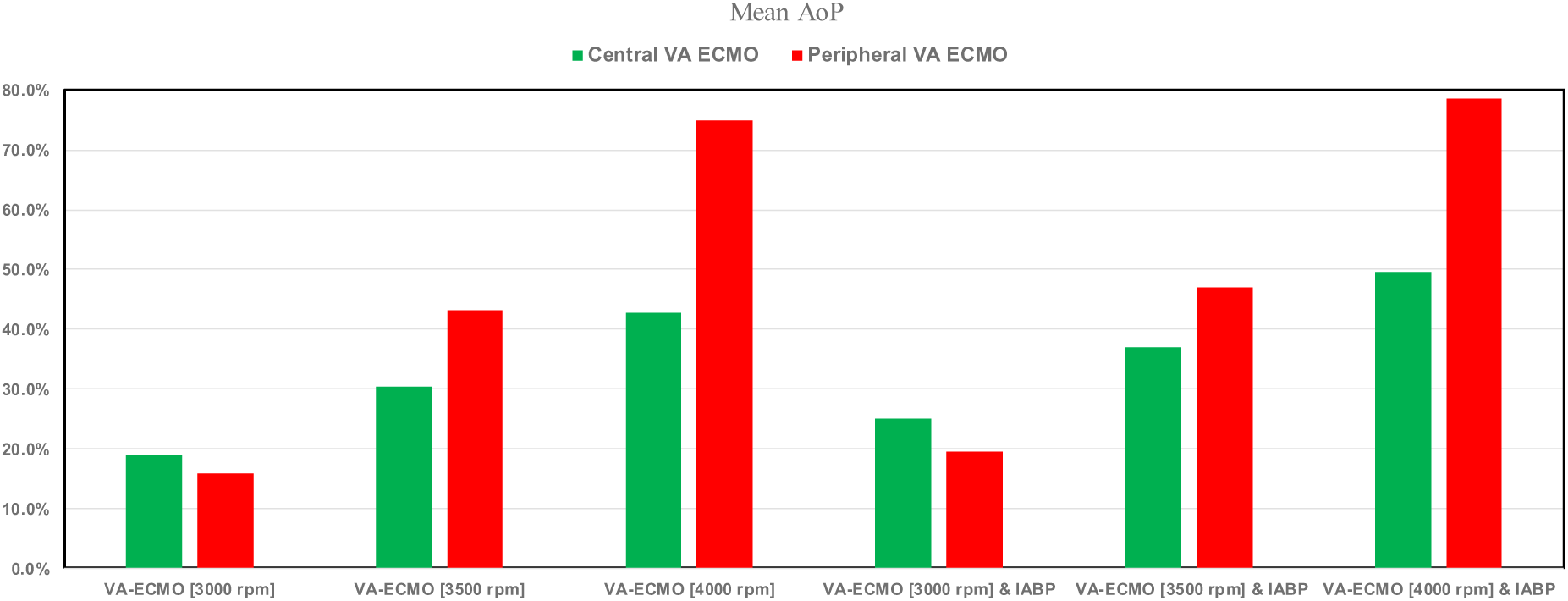
Percentage changes in mean AoP value calculated in relation to pathological conditions for both central and peripheral VA ECMO assistance (with pump rotational speed set to 3000, 3500, and 4000 rpm), with and without IABP support.

The effect caused by central (top panel) and peripheral (bottom panel) VA ECMO support and IABP assistance on mean wedge and mean pulmonary arterial pressures and on minimum and maximum pulmonary arterial pressure is shown in Figure 10.

**Figure 10.**
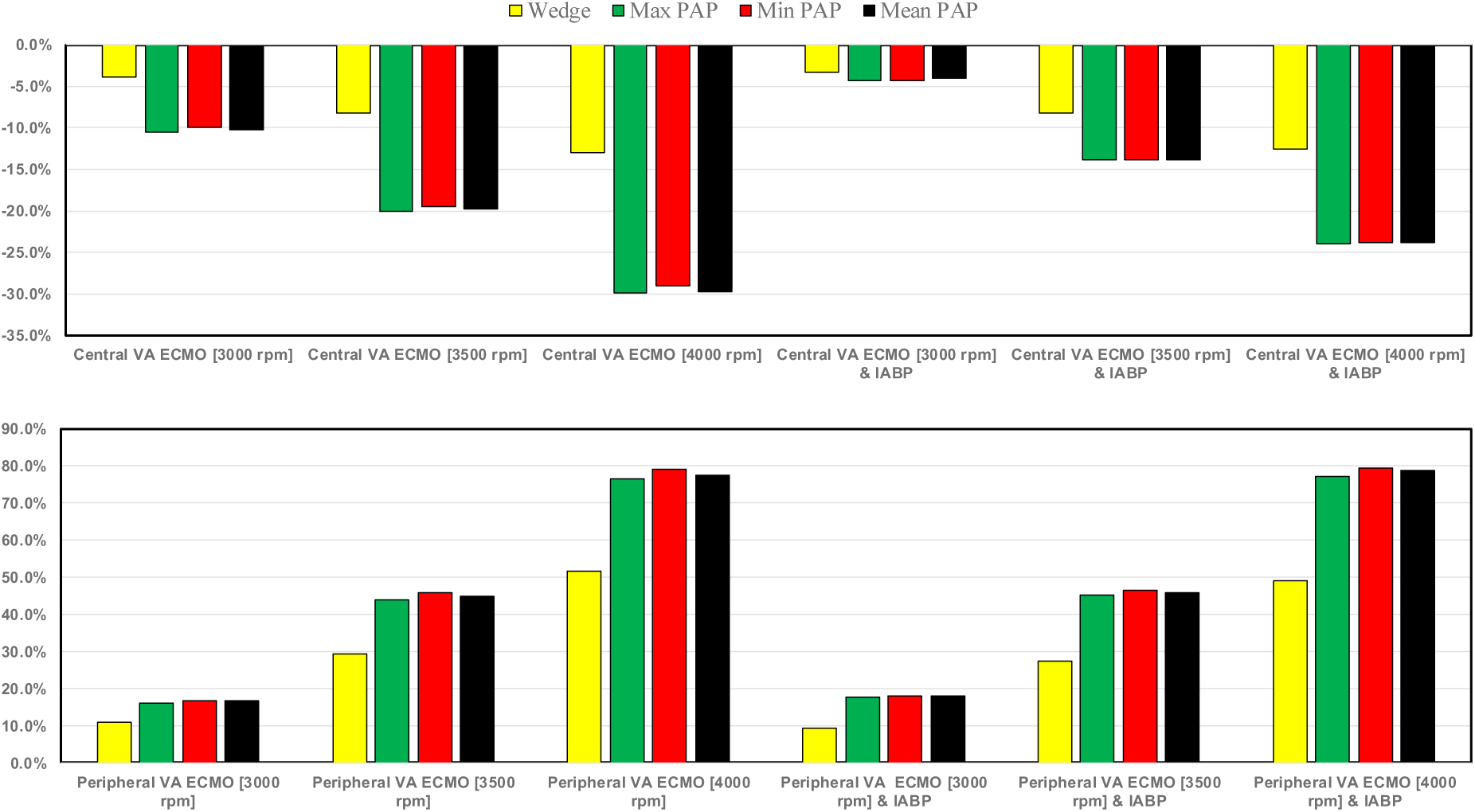
Percentage changes in mean wedge value and minimum, mean and maximum pulmonary arterial pressure values calculated in relation to pathological conditions for both central and peripheral VA ECMO assistance (with pump rotational speed set to 3000, 3500, and 4000 rpm), with and without IABP support.

When VA ECMO was configured in central mode with high pump rotational speeds, the wedge drastically reduced; under these circumstances, IABP support appeared to have little effect on the wedge (top panel). Peripheral VA ECMO increased the wedge by approximately 50% (compared to pathological conditions) at a pump speed of 4000 rpm (bottom panel). The minimum, maximum, and mean pulmonary arterial pressures decreased up to 30% for central VA ECMO (with a pump rotational speed of 4000 rpm); when IABP and VA ECMO were activated simultaneously, the reduction was less (top panel). Peripheral VA ECMO with high pump rotational speed (4000 rpm) increased the minimum, mean and maximum values of pulmonary arterial pressure up to 80%. These effects did not appear to be influenced by the presence of IABP. Figure 11 shows that central VA ECMO support reduced the right atrial pressure-volume loop area from 15% to 50.7% without IABP assistance (top panel). IABP activation with central VA ECMO led to the reduction in RA-PVLA by a maximum value of 38.3% (when pump rotational speed was set to 4000 rpm).

**Figure 11.**
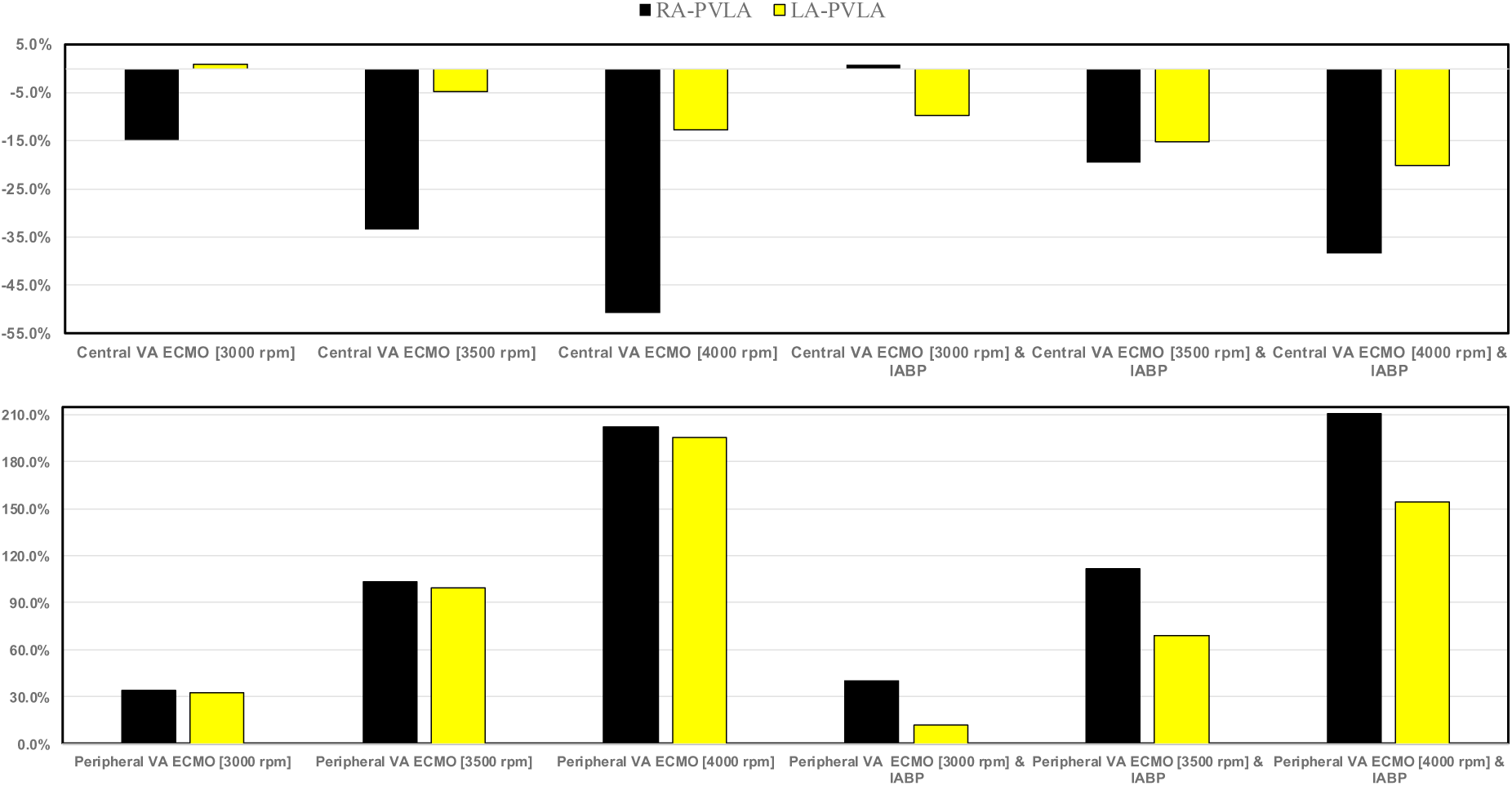
Percentage changes in right and left atrial pressure-volume loop area in relation to pathological conditions for both central (top panel) and peripheral (bottom panel) VA ECMO assistance (with pump rotational speed set to 3000, 3500, and 4000 rpm), with and without IABP support.

When simulating double assistance, the percentage decrease in comparison to the pathological state of LA-PVLA was more noticeable. Peripheral VA ECMO increased both RA-PVLA and LA-PVLA when the pump rotational speed increased, as seen in the bottom panel (Figure 11). Our simulations indicate that a minor increase in LA-PVLA was induced by IABP support.

The results of the simulation for LVEDV, LVESV, RVEDV and RVESV are shown in Figure 12. The top (bottom) panel shows the percentage changes under assisted conditions with central (peripheral) VA ECMO driven at 3000, 3500, and 4000 rpm, as well as with and without IABP support, related to pathological conditions. When IABP assistance was not active, central VA ECMO (top panel) increased LVESV (LVEDV) from 12.7% to 26.7% (from 6.1% to 10.7%) whilst decreased RVESV (RVEDV) from 3.9% to 14.1% (from 7.5 to 22.7%). IABP activation caused a percentage drop in LVEDV, RVESV and RVEDV for each pump rotational speed, whereas LVESV increased in percentage as the pump speed increased.

**Figure 12.**
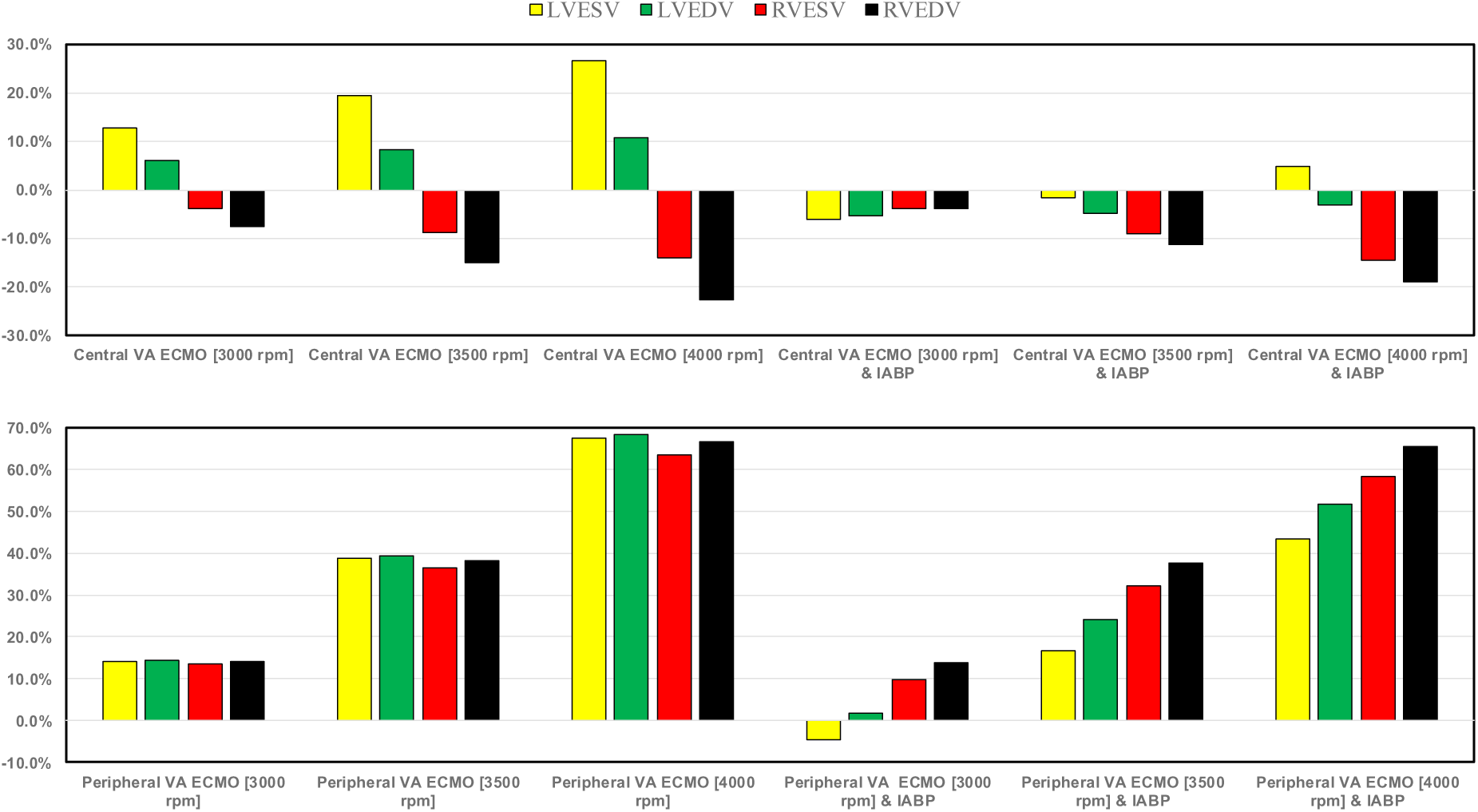
The top (bottom) bar graph shows the percentage changes in LVESV, LVEDV, RVESV and RVEDV under assisted conditions with central (peripheral) VA ECMO driven at 3000, 3500, and 4000 rpm, as well as with and without IABP support, in relation to pathological conditions.

Peripheral VA ECMO (bottom panel) caused a similar percentage increase in LVESV, LVEDV, RVESV and RVEDV at each pump speed (e.g., around 14% at 3000 rpm). IABP activation produced different percentage change increases in the four parameters (e.g., 16.7% in LVESV, 24.0% in LVEDV, 32.3% in RVESV, and 37.8% in RVEDV at 3500 rpm, respectively).

Figure 13 shows the effects induced on the energetic variables by central (peripheral) VA ECMO with and without IABP activation. IABP support combined with central VA ECMO reduced LVEW, RVEW, and RPVVA (top panel) at all pump rotational speeds. When the pump rotational speed was set to 4000 rpm, the reduction in percentage changes in LVPVA was negligible, whereas RVEW, RVPA, and LVEW decreased by 39.5%, 37.6%, and 15.6%, respectively. Simulations performed with only central VA ECMO assistance increased LVPVA up to 39.2% (pump rotational speed of 4000 rpm), and reduced RVEW and RVPVA up to 48.8% and 44.8%, respectively (pump rotational speed of 4000 rpm). LVEW, LVPVA, RVEW, and RVPVA significantly increased in simulations with only peripheral VA ECMO support (bottom panel); this effect became more evident as the pump rotational speed increased. While RVEW and RVPVA seemed to be unaffected by the presence of IABP, concomitant IABP support decreased LVEW and LVPVA for all values of pump rotational speed.

**Figure 13.**
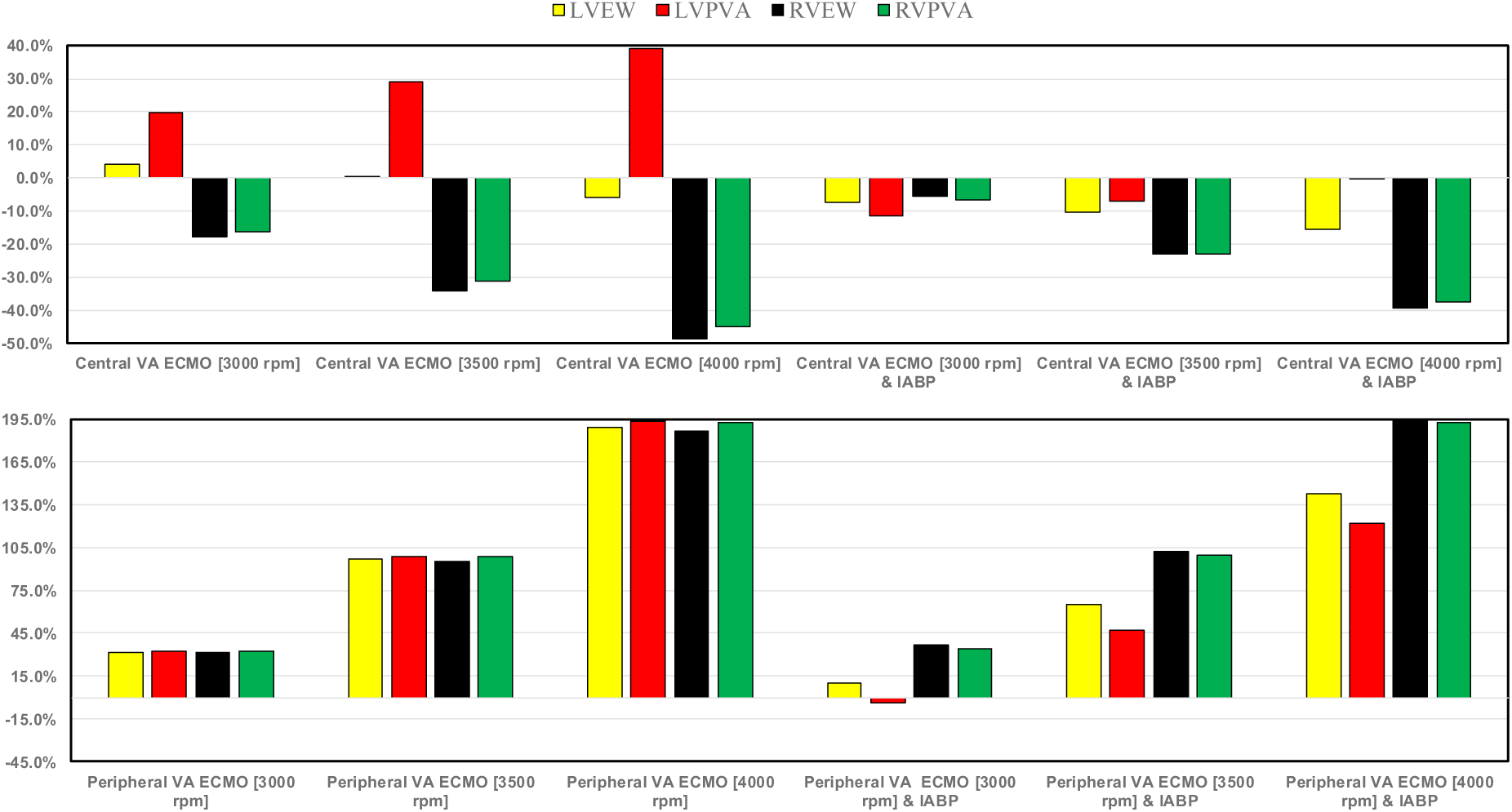
The top (bottom) panel shows the percentage changes in LVEW, LVPVA, RVEW and right ventricular pressure-volume area under assisted conditions with central (peripheral) VA ECMO driven at 3000, 3500, and 4000 rpm, as well as with and without IABP support, in relation to pathological conditions.

## Discussion

The use of VA ECMO support in critically ill patients remains a widely recognised approach despite current controversy about its beneficial effects and outcomes. The use of IABP to address LV unloading has been an appropriate and recognised course of action. Nevertheless, there remains additional controversy related to its use with either peripheral or central VA ECMO support. This has given us the motivation to review current views and shed some lights on the subject by answering the following question: is there a real difference between central and peripheral VA ECMO when associated with IABP. We sought to address this question in the context of a simulation setting based on lumped parameter modelling and pressure-volume analysis. Initial simulations with pump rotational speed at 3000 rpm revealed that central VA ECMO support caused less LV and RV distension compared to peripheral VA ECMO with reduction of RVESV, RVEDV and LAEDV as shown in Figure 3. When combined IABP and VA ECMO support was activated, central VA ECMO clearly unloaded both ventricles compared to peripheral VA ECMO as shown in Figure 4 and Figure 12. Central VA ECMO at 3000 and 3500 rpm would increase SVRi and decrease CI whilst peripheral VA ECMO would increase CI without any effect on SVRi. The concomitant use of IABP for both configurations did not make any significant difference. Central VA ECMO at 3000 and 3500 rpm increased systemic Ea up to 40% and Ea/Ees ratio up to 52%. The combined use of central VA ECMO and IABP would increase both parameters by only 15% (Figure 5). Peripheral VA ECMO at 3000 and 3500 rpm had a negligible effect on systemic Ea and Ea/Ees ratio whilst the combined use with IABP decreased both parameters by 20%. The effect of both VA ECMO configurations with or without IABP was negligible on pulmonary Ea (Figure 5). The Ea/Ees ratio reflects effective ventricular-arterial coupling, which can be defined as the provision of optimal cardiovascular flow reserve without compromising arterial blood pressure. Central VA ECMO alone seems to induce a more pronounced deleterious effect compared to peripheral VA ECMO. Arguably, the combined use of central VA ECMO and IABP seems to largely mitigate this negative effect. Central VA ECMO increased coronary blood flow (CBF) up to 55.82% with pump rotational speed at 4000 rpm. IABP addition would increase CBF up to 61.3% (Figure 6). Peripheral VA ECMO increased CBF up to 65.15% with the same rotational pump speed. The combined use with IABP would increase CBF up to 68.66% (Figure 6). The effect on cerebral blood flow appeared more pronounced under central VA ECMO support with low rotational speed (Figure 7). The use of peripheral VA ECMO alone or combined with IABP would increase left atrial pressure (LAP), left ventricular pressure (LVP) and aortic pressure (AoP). In contrast, central VA ECMO alone would not increase these parameters as much. The combined use of IABP would either reduce these parameters or increase them to a lesser extent (Figure 8 and Figure 9). Central VA ECMO alone would reduce pulmonary capillary wedge pressure (PCWP) and mean pulmonary artery pressure (PAP). The concomitant use of IABP would maintain this trend (Figure 10). Peripheral VA ECMO alone or combined with IABP would generate the opposite effect on PCWP and mean PAP (Figure 10).

From an energetic point of view, central VA ECMO alone or combined with IABP has a much more favourable profile compared to peripheral VA ECMO alone or combined with IABP (Figure 11 and Figure 13). Although peripheral VA ECMO has been used more frequently even in postoperative cardiac patients (Chen, 2019; Baran, 2024; Nishi, 2022; Li, 2019), myocardial and cerebral blood flow remain negatively affected (Bělohlávek, 2012; Yang, 2014; Xu, 2014). In contrast, the outcome of our simulations has revealed a favourable effect on coronary blood flow for both peripheral and central VA ECMO at 4000 rpm pump rotational speed, either alone or combined with IABP. A pump rotational speed at 3000 rpm would lead to a more pronounced effect on CBF under central VA ECMO alone or combined with IABP compared to peripheral VA ECMO alone or combined with IABP. A similar trend was observed for the cerebral blood flow under central VA ECMO support at 3000 rpm pump rotational speed, either alone or combined with IABP. Nevertheless, the overall haemodynamic and energetic analysis of our simulations support the use of central VA ECMO and IABP in line with previous experimental work (Sauren, 2007; Farag, 2022; Gerfer, 2023; Djordjevic, 2022; Petroni, 2014). We would not preclude the use of peripheral VA ECMO combined with IABP although central VA ECMO would be advisable in postcardiotomy failure patients or whenever feasible.

## Conclusion

VA ECMO can provide effective circulatory support in critically ill patients due to cardiogenic shock. IABP is appropriate to unload the left ventricle and counteract the detrimental effect of full VA ECMO support. A combined use of central VA ECMO and IABP may be the preferred approach when feasible.

## Notes

### Competing Interest Statement

The authors have declared no competing interest.

